# Tetrasodium EDTA disrupts *Pseudomonas aeruginosa* membrane integrity, shows suppressed resistance evolution and reduced cytotoxicity compared to meropenem

**DOI:** 10.64898/2026.08.11.744140

**Authors:** Oluwatosin Qawiyy Orababa, Charlotte Cornbill, Ayomikun Kade, Chidiebere F. Uchechukwu, Shristi Sharma, Leonard Uzaiure, Natasha Reddy, Rupika Gulati, Blessing Oyedemi, Freya Harrison

## Abstract

*Pseudomonas aeruginosa* remains one of the most important clinical pathogens for which new drugs are needed, due to its resistance machinery. Consequently, there is an increasing effort to develop new and effective treatments against this pathogen. We recently showed that tetrasodium ethylenediaminetetraacetic acid (tEDTA) exhibits promising antibacterial and antibiofilm activity against *P. aeruginosa* in advanced biofilm models. tEDTA is known to chelate divalent cations, with predicted effects on the outer membrane; however, a full understanding of how this kills *P. aeruginosa* is lacking. Also, it is currently not clear how slowly or rapidly *P. aeruginosa* will evolve resistance to this treatment. Using membrane disruption assays and RNA-seq, we showed that tEDTA disrupts bacterial membrane potential and permeabilises *P. aeruginosa* membranes. RNA-seq revealed the significant upregulation of genes involved in the transport of iron, phosphate, potassium, and magnesium ion. The *arnABCD* operon which is involved in lipid A biosynthesis was also upregulated. Using a 7-day evolutionary ramp approach, we showed that *P. aeruginosa* could not evolve resistance to tEDTA under strong selection. Lastly, we carried out a cytotoxicity assay with Human Epithelial type 2 (HEp-2) cells and showed that there was reduced cytotoxicity of tEDTA compared to meropenem. This study provides good insight into the mechanism of action of tEDTA and further evidence of its potential as an alternative to antibiotics for *P. aeruginosa* infections.

## INTRODUCTION

*Pseudomonas aeruginosa* is a Gram-negative opportunistic bacterium, notorious for its resistance to a broad array of antibiotics (Elfadadny et al., 2024). In an escalating crisis of antimicrobial resistance (AMR), *P. aeruginosa* poses one of the most significant threats to global public health, undermining decades of progress in the treatment of infectious diseases (Miller and Arias, 2024). As a leading cause of healthcare-associated infections, including ventilator-associated pneumonia, catheter associated urinary tract infections, and wound infections, *P. aeruginosa* commonly exhibits multidrug-resistant (MDR) and even extensively drug-resistant (XDR) phenotypes (Coello-Pelegrin et al., 2021; Sathe et al., 2023). These phenotypes have led to *P. aeruginosa* being classified by the World Health Organization as a high-priority pathogen, for which the development of new treatment options is needed (Alharbi et al., 2025). There is a focus on antibiotic alternatives that can disarm resistance mechanisms by disrupting biofilm matrix, permeating through the biofilm matrix to reach cells, and/or overcoming the Gram-negative outer membrane barrier.

One potential antibiotic alternative that has regained a renewed interest is tetrasodium ethylenediaminetetraacetic acid (tEDTA). The antibacterial and antibiofilm potential of tEDTA have previously been reported in different studies (Liu et al., 2018; Percival and Salisbury, 2018; Crowther et al., 2025; Orababa et al., 2026). Moreover, a 4% (wt/vol) solution of tEDTA has been approved for use as a lock solution to prevent biofilm colonisation and blockage of vascular catheters in Canada (Liu et al., 2018). In our recent study we showed that the antibacterial activity of tEDTA can be replicated in different growth environments, including high-validity models of cystic fibrosis lung, chronic wound and endotracheal tube biofilm (Orababa et al., 2026). We also reported that 1-2% wt/vol tEDTA (below the clinically approved concentration) retains potent antibiofilm efficacy against *P. aeruginosa* grown in clinically relevant models of chronic wounds and cystic fibrosis (Orababa et al., 2026), which are rich in divalent cations and cue higher levels of tolerance to antibiotics and antimicrobials than simpler high-throughput platforms (Werthén et al., 2010; Sweeney et al., 2020; Walsh et al., 2024).

Currently, the mechanism of action and molecular target(s) of tEDTA in *P. aeruginosa* is not fully understood. The fundamentals behind EDTA’s ability to affect *P. aeruginosa* growth stem from its ability to chelate divalent metal cations, specifically magnesium (Mg²) and calcium (Ca²) (Banin et al., 2006). A more detailed understanding of how tEDTA affects *P. aeruginosa* cells, and how *P. aeruginosa* responds to tEDTA on a cellular (tolerance) and evolutionary (resistance) scale is critical for the investigation and development of tEDTA as an antimicrobial. This information will inform strategies such as combination treatment with antibiotics to achieve synergy, or optimisation of treatment protocols to mitigate or circumvent resistance evolution (concentrations or repeated dosing).

Further, EDTA’s potent calcium chelation properties could raise concerns for systemic toxicity, such as hypocalcaemia or cardiac effects, which could limit tEDTA’s use to topical and local application (e.g. catheter lock solutions or wound care products (Finnegan and Percival, 2015). Hence, understanding its effects on human cells is essential in its clinical application, especially as a topical agent in skin and soft tissue infections. Currently, no study has assessed the cytotoxicity levels of tEDTA on human epithelial cells. This is important as we have recently shown tEDTA to have good antibiofilm efficacy against *P. aeruginosa* in a soft tissue biofilm model, evidencing its potential in wound care (Orababa et al., 2026).

In this study, we report the antibacterial efficacy of tEDTA against both standard laboratory and clinical strains of *P. aeruginosa*. We also showed that the antibacterial activity of tEDTA was due to its ability to cause significant disruption of both the outer and inner membrane of *P. aeruginosa*. For the first time, we revealed the upstream and downstream effects of tEDTA on *P. aeruginosa* and provided more detailed understanding of its antibacterial effect. The ability of bacteria to rapidly evolve resistance to antimicrobials is one of the biggest challenges of the twenty-first century. Pathogens like *P. aeruginosa* are notorious for this, due to their arsenal of both intrinsic and acquired resistance mechanisms. For the first time, we show that it is more difficult for *P. aeruginosa* strains to evolve resistance to tEDTA than to meropenem under lab conditions that impose strong selection for resistance.

## RESULTS

### Tetrasodium EDTA has strong antibacterial efficacy against both standard laboratory and clinical strains of *P. aeruginosa*

We previously showed that tEDTA has good antibacterial efficacy against the *P. aeruginosa* lab strain and strong biofilm former PA14 (Orababa et al., 2026). We extended our assessment to a wider strain panel including the lab strains PAO1 and LESB58 and seven clinical isolates derived from cystic fibrosis lung infection, as different strains may respond differently to the same antimicrobial. We used % wt/vol to measure tEDTA minimum inhibitory concentration (MIC), using a broth microdilution assay in cation-adjusted Muller-Hinton broth (caMHB). In parallel, we have also provided the antimicrobial susceptibility profile of the clinical strains to clinically-relevant antibiotics (meropenem, tobramycin, and ciprofloxacin). We classified good susceptibility to tEDTA as MICs below 1 % wt/vol of tEDTA. We observed a consistent tEDTA MIC of 0.25% across the three laboratory strains, which included one strain classed as clinically resistant to meropenem (LESB58) (Table 1). Some of the clinical strains had tEDTA MIC values below 0.25% and their tEDTA susceptibility levels corresponded to their meropenem susceptibility levels. Strain PA30 with meropenem resistant profile had the highest tEDTA MIC (0.25%) among the clinical strains while the lowest tEDTA MIC was observed in PA68. Interestingly, we also report good susceptibility to tEDTA among multidrug-resistant clinical strains of *P. aeruginosa* (PA1 H142500397 and PA2 H142340666).

**Table 1.** Minimum inhibitory concentration of tEDTA against both standard laboratory strains and clinical isolates of *Pseudomonas aeruginosa*. Antimicrobial susceptibility profiles of *P. aeruginosa* strains were determined with broth microdilution or disc diffusion methods. Strains were classified as resistant (R), intermediate (I), or susceptible (S) to antibiotics based on EUCAST recommendation

| *P. aeruginosa* Strain | Type | MIC of treatments, or result of disk diffusion assay |  |
| --- | --- | --- | --- |
|  |  | tEDTA (% wt/vol) | Antibiotics |
| PA14 | Laboratory strain | 0.25% (6.5 mM) | Meropenem -1 µg/mL<sup>S</sup> (3.0 µM) |
| LESB58 | Laboratory strain | 0.25% (6.5 mM) | Meropenem -16 µg/mL<sup>R</sup> (47.9 µM) |
| PAO1 | Laboratory strain | 0.25% (3.3 mM) | N/A |
| PA66 | Clinical strain | 0.125% (6.5 mM) | Meropenem - 4 µg/mL<sup>I</sup> (12.0 µM) |
| PA30 | Clinical strain | 0.25% (6.5 mM) | Meropenem - 16 µg/mL<sup>R</sup> (47.9 µM) |
| PA21 | Clinical strain | 0.125% (3.3 mM) | Meropenem - 8 µg/mL<sup>I</sup> (23.9 µM) |
| PA68 | Clinical strain | 0.063% (1.6 mM) | Meropenem - 0.5 µg/mL<sup>S</sup> (1.5 µM) |
| PA1 H142500397 | Clinical strain | 0.063% (1.6 mM) | * Tobramycin (R), Meropenem (R),<br>Ciprofloxacin (R) |
| PA2 H142340666 | Clinical strain | 0.125% (3.3 mM) | * Tobramycin (R), Meropenem (R),<br>Ciprofloxacin (R) |
| PA3 H142520850 | Clinical strain | 0.125% (3.3 mM) | * Tobramycin (R), Meropenem (S),<br>Ciprofloxacin (S) |

\*Susceptibility profile done with disk diffusion method
<sup>S</sup> represents susceptible, <sup>I</sup> represents intermediate and <sup>R</sup> represents resistant, according to the EUCAST breakpoint for meropenem (EUCAST, 2024).

To understand if susceptibility/resistance profile of the strains to meropenem influence the bacteriostatic effect of tEDTA on *P. aeruginosa* strains, we assessed the planktonic growth kinetics of PA14 and LESB58 in the presence of 0.25-4×MIC tEDTA in caMHB. These strains were chosen to represent meropenem-susceptible and resistant strains of *P. aeruginosa*, respectively. We observed some differences in the growth kinetics of these two strains as PA14 grew to a higher biomass (OD_600_) compared to LESB58 in the untreated wells (Fig 1). Interestingly, a slight increase in OD_600_ was observed at 1×MIC of tEDTA with LESB58 but not with PA14 indicating a slight tolerance to tEDTA, however, this change was not detectable by eye (Fig 1).

**Fig 1.**
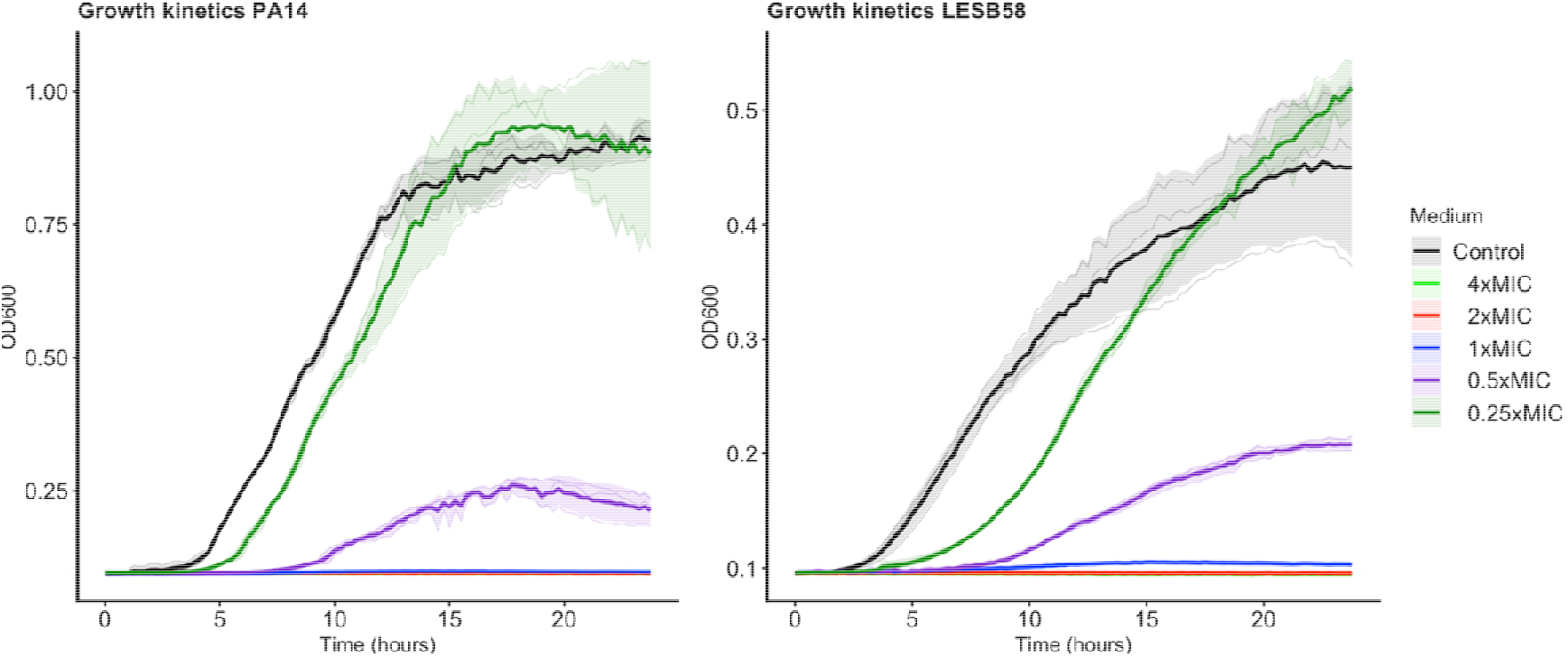
Growth kinetics of *P. aeruginosa* PA14 and LESB58 upon exposure to different concentrations of tEDTA. Overnight cultures of *P. aeruginosa* PA14 and LESB58 were diluted to 0.01-0.008 OD_600_ and treated with different concentrations of tEDTA ranging from 0.25×MIC to 4×MIC. These were incubated in a TECAN Spark multimode plate reader at 37°C for 24 h and OD_600_ was measured every 15 minutes. Results shown are from three biological replicates; each of the replicates is shown as an individual line, with the mean and standard deviation plotted as a heavier line and shaded interval.

### tEDTA disrupts bacterial membrane integrity

Despite different reports of the antibacterial activity of tEDTA (Liu et al., 2018; Crowther et al., 2025; Orababa et al., 2026), its mechanism of action has not been well elucidated, especially against *P. aeruginosa*. Moore et al. (2023) previously reported that tEDTA has high cation-chelating activity, similar to EDTA. Bacterial membranes (inner and outer membrane) contain cations that play an essential role in their stability (Clifton et al., 2015; Bashford et al., 1988), hence, chelation of these ions can cause significant disruption of bacterial membrane integrity and subsequently lead to growth inhibition/death.

To understand the effect of tEDTA on bacterial membrane, we used the membrane-sensitive dye DiSC3(5). When DiSC3(5) is added to bacteria with stable membrane potential, it binds to the plasma membrane, leading to reduced fluorescence of DiSC3(5). When a compound that has the ability to disrupt bacterial inner membrane is added, the fluorescence of DiSC3(5) is either rapidly decreased or increased indicating membrane permeabilization (depolarization) or membrane potential disruption (hyperpolarization) respectively. A 0.6% wt/vol (above MIC) concentration of tEDTA was used for this assay to allow clearer observation of membrane disruption due to the higher cell concentration (0.5 OD_600_) used for this assay compared to the MIC and growth kinetics assays (1 in 100 of 0.1 OD_600_). Upon the addition of 0.6% tEDTA, we observed a rapid and strong decrease in the fluorescence of DiSC3(5), similar to what we observed with CCCP – a known bacterial membrane disruptor (Fig 2a). This evidences the ability of tEDTA to cause observable disruption of bacterial membrane. Initially, we observed a rapid quenching of DiSC3 upon the addition of tEDTA which was then followed by a slow dequenching (Fig 2a). This suggests an initial rapid permeabilization of the outer membrane followed by a slow depolarisation of the inner membrane. This is similar to the previously reported dual-membrane disruptive activity of the well-characterized antibiotic polymyxin B (Buttress et al., 2022).

**Figure 2.**
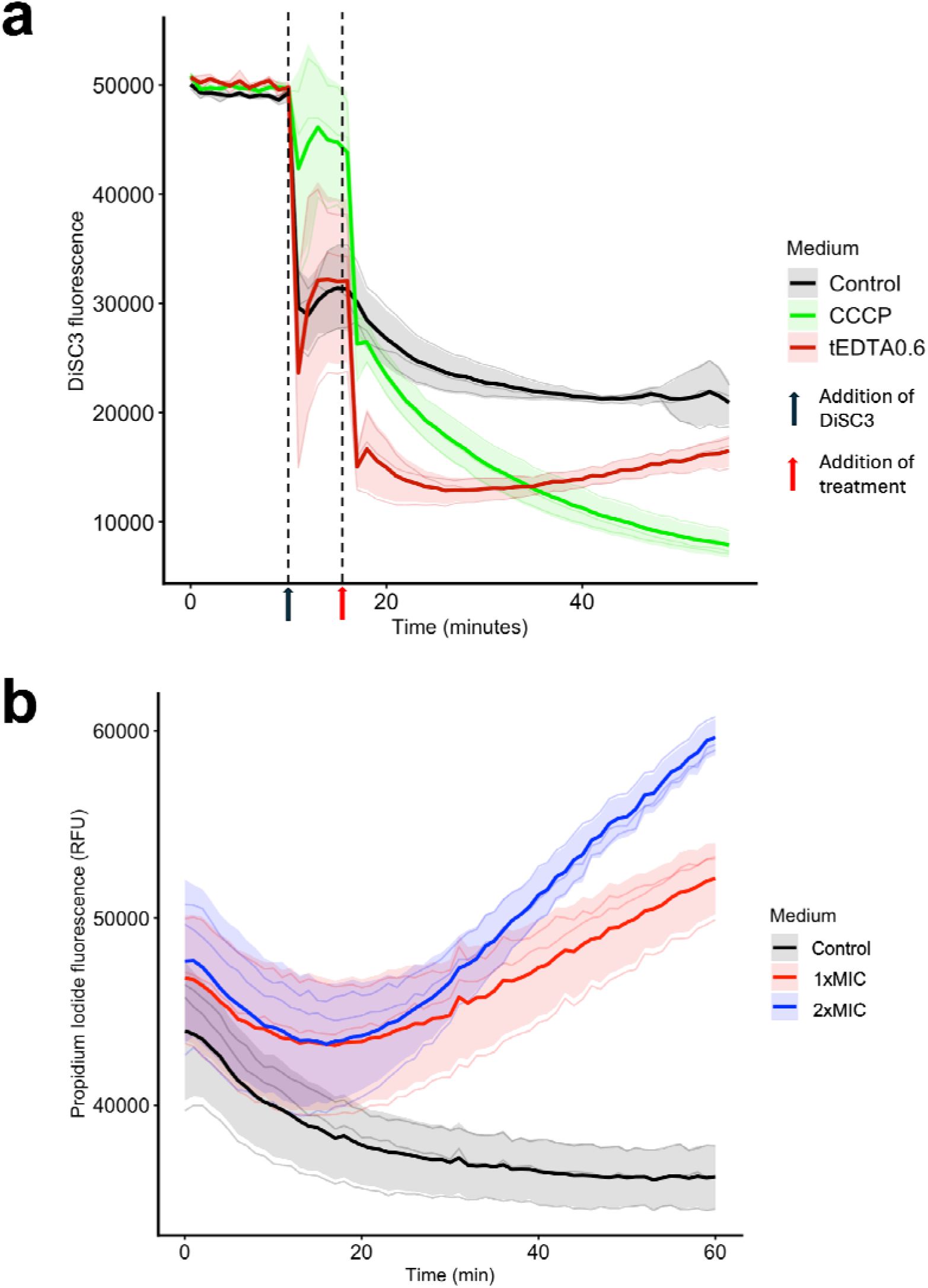
tEDTA permeabilises and depolarises *P. aeruginosa* membranes. **a.** *P. aeruginosa* (0.5 OD_600_) was stained with DiSC3(5), treated with 0.6% tEDTA (red), 20 µg/ml CCCP (green), or water (black), and their fluorescence monitored for 1 h. There was a sharp decrease in DiSC3(5) fluorescence when tEDTA or CCCP were added, indicating membrane disruption. The red arrow indicates point of treatment. Results shown are from three biological replicates; with the mean and standard deviation plotted as a heavier line and shaded interval. **b.** *P. aeruginosa* (0.1 OD_600_) was treated with 1×MIC (red), 2×MIC of tEDTA (blue), or water (black) and stained with 5 µg/ml propidium iodide, and their fluorescence monitored for 1 h. Increase in propidium iodide fluorescence indicate. The red arrow indicates point of treatment. Results shown are from three biological replicates; with the mean and standard deviation plotted as a heavier line and shaded interval.

To further confirm if there was permeabilization of the *P. aeruginosa* inner membrane by tEDTA, we carried out a propidium iodide assay. Propidium iodide is a membrane impermeable nucleic acid dye whose fluorescence increases when bound to the nucleic acid of permeabilised cells. This assay confirms membrane permeabilization as we observed an increase fluorescence of propidium iodide in the tEDTA treated cells. This was not seen in the untreated cells (Fig 2b).

To further confirm the disruption of membrane integrity, we also carried out bacterial cytological profiling with Sytox green, FM4-64, and DAPI. Sytox green is another membrane impermeable dye whose fluorescence significantly increases when bound to the DNA of a permeabilised cell while DAPI is a membrane permeable dye that stains both permeabilised and impermeabilized cells. FM4-64 stains the bacterial cytoplasmic membrane and its fluorescence increases when the cytoplasmic membrane integrity is disrupted. There was increased fluorescence of both FM4-64 and Sytox green, showing evidence of outer and inner membrane disruption by tEDTA (Fig 3; Supplementary File).

**Fig 3.**
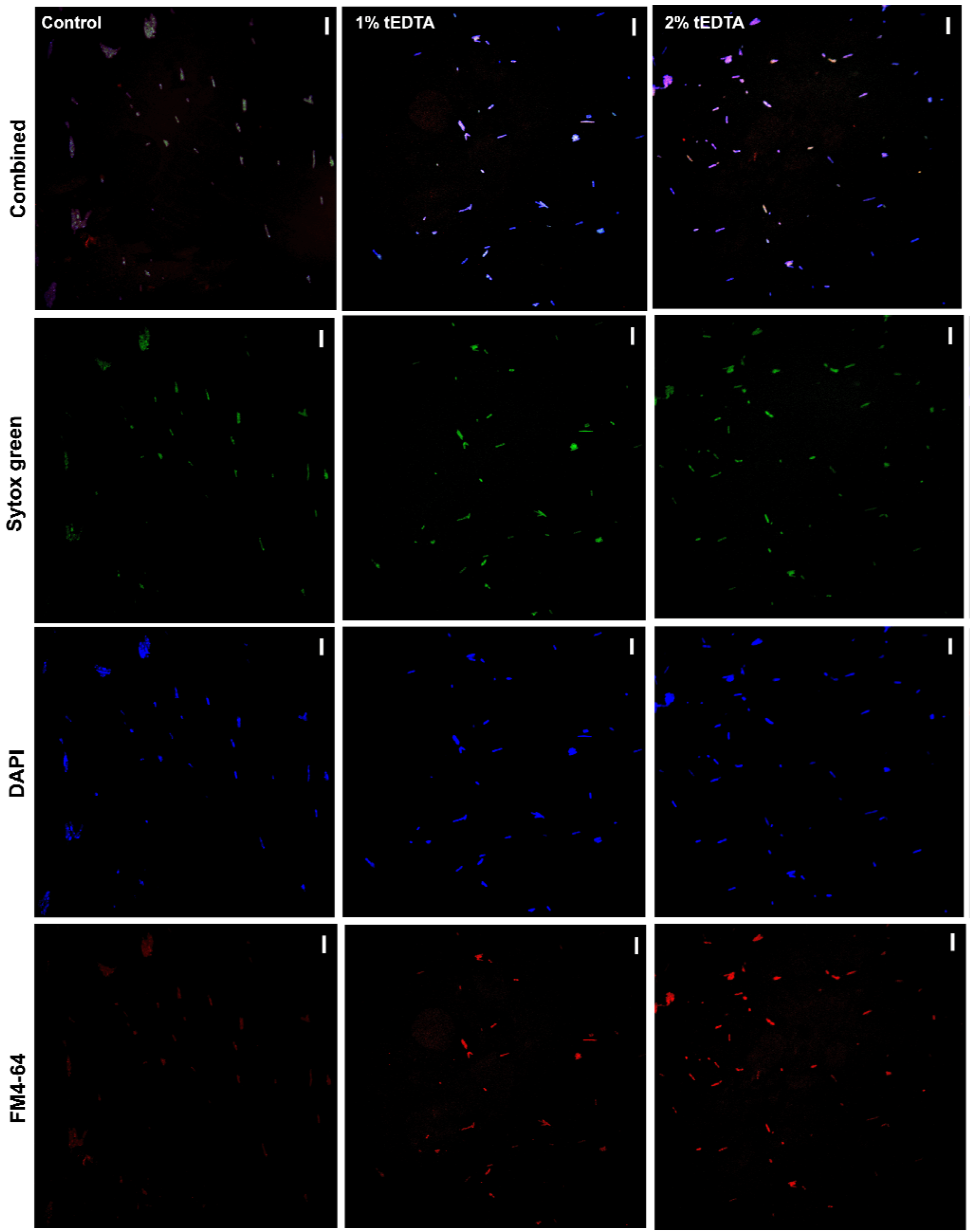
Bacterial cytological profiling of *P. aeruginosa* treated with tEDTA. *P. aeruginosa* was treated with 1% wt/vol (4×MIC) tEDTA, 2% wt/vol (8×MIC) tEDTA or sterile water (control) and stained with SYTOX green, DAPI, and FM4-64. The cells were then imaged with 100x oil immersion objective on a Nikon Widefield Ti2-E (NW1) microscope.

These results evidence membrane potential disruption and membrane permeabilization by tEDTA.

### Transcriptomics data reveals increased expression of genes associated with membrane transport and siderophore activities

Bacterial membrane potential plays an important role in bacterial physiological functions such as transmembrane transport, bacterial chemotaxis, and energy synthesis (Benarroch and Asally, 2020). Having previously shown in Fig 2 and Fig 3 that tEDTA disrupts bacterial membrane integrity via the disruption of bacterial membrane potential and membrane permeabilistion, we wanted to further understand the downstream effect of this disruption on *P. aeruginosa* physiological responses. We assessed this by looking at the *P. aeruginosa* gene expression in response to tEDTA using RNA-seq. Mid-exponential phase populations of PA14 were treated with 4× MIC tEDTA (1% wt/vol) for one hour, or left untreated. Again, a high concentration of tEDTA was used here due to the high cell population used (final cell population of 0.4 OD_600_) in this assay and to allow effects of tEDTA to be observed before cells begin to die. RNA was extracted and mRNA sequenced.

Functional annotation analysis of differentially expressed genes (DEGs: |Log_2_foldchange| ≥ 1.5 and p_adj_ < 0.05) revealed the significant upregulation of Kyoto Encyclopedia of Genes and Genomes (KEGG) groups associated with biosynthesis of siderophore group non-ribosomal peptides, two-component systems, biosynthesis of secondary metabolites, ABC transporters and bacterial chemotaxis among others (Fig 4). Similarly, Gene Ontology (GO) groups associated with ion transport, response to stimulus, cell communication and locomotion, among others, were significantly upregulated (Fig 4) The top 50 upregulated and downregulated GOs with their corresponding genes have been included in Supplementary File 1.

**Fig 4.**
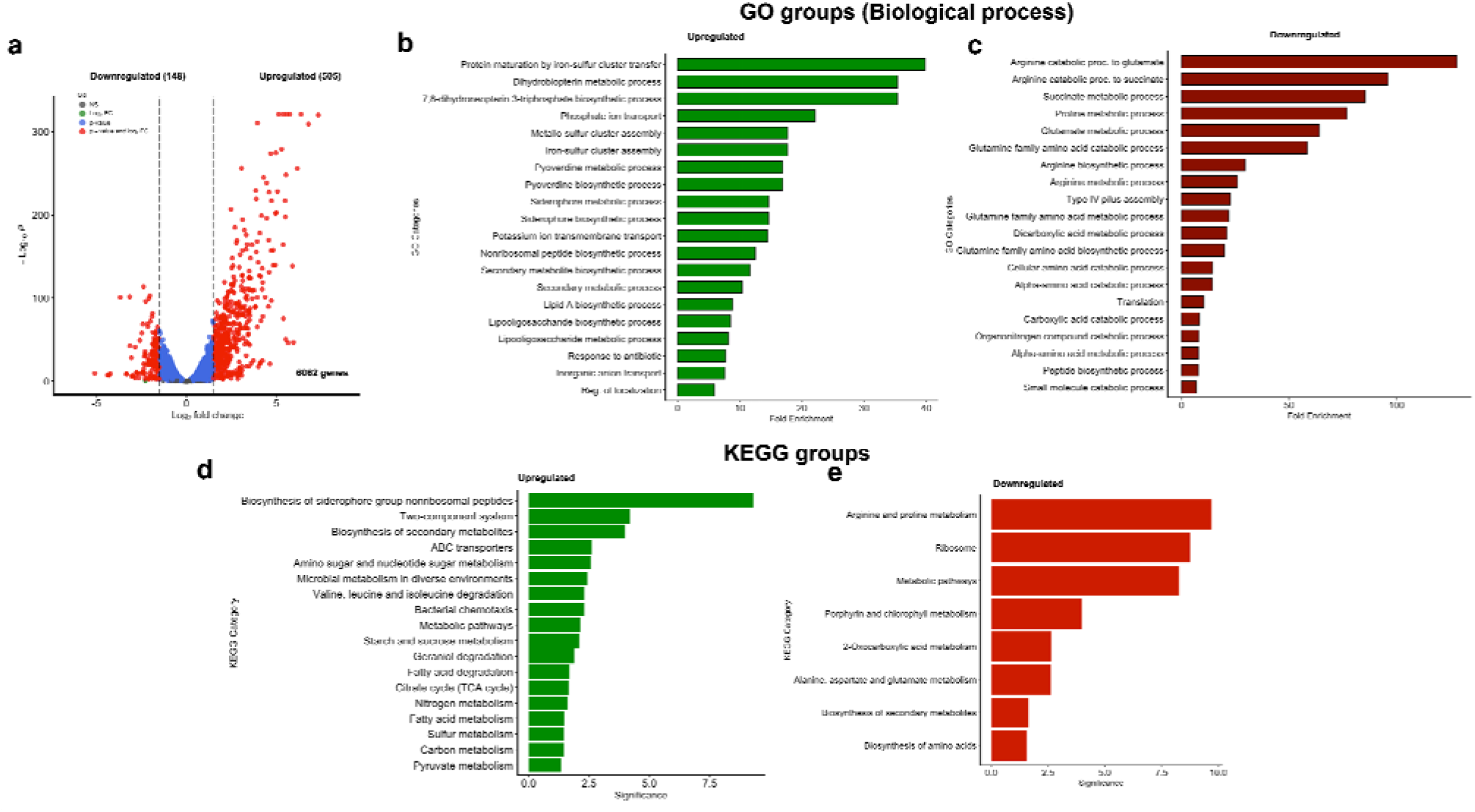
Significantly enriched GO and KEGG groups in *P. aeruginosa* PA14 in response to 4×MIC of tEDTA. *P. aeruginosa* cells at mid-exponential phase were treated with 4× MIC (1% wt/vol) of tEDTA for 1 h before RNA sequencing. The differentially expressed genes (p-values ≤ 0.05 and |log_2_foldchange| ≥1.5) in the treated vs untreated cells were analysed (**a**). The top 20 upregulated (green) and downregulated (red) gene ontology (GO) groups were also identified (**b, c**). **d** and **e** are KEGG groups from upregulated and downregulated genes, respectively.

Some of the genes that were differentially upregulated in response to tEDTA include *pstA*, *pstB*, *pstS*, *phoU*, *phoB*, *kdpA*, *kdpB*, *kdpC*, *mgtA*, *cntO*, *pvdQ*, *pvdA*, *pvdM*, *pvdN*, *pvdE*, and *pvdF* (Fig 5). Due to the roles of these transmembrane ion transport genes in ion (phosphate, potassium, magnesium, and iron) uptake, their upregulation is an indication of ion limitation. Fig 5 is a schematic representation of upregulated genes, and their log_2_foldchange values, associated with ion transport and the role of their product in transmembrane ion transport. GO groups associated with lipid A and lipopolysaccharide biosynthesis were also enriched and the genes in this group are the *arnABCD* genes (Fig 3). This further evidence the effect of tEDTA on the Gram-negative membrane. Altogether, these confirm the membrane activity of tEDTA.

**Fig 5.**
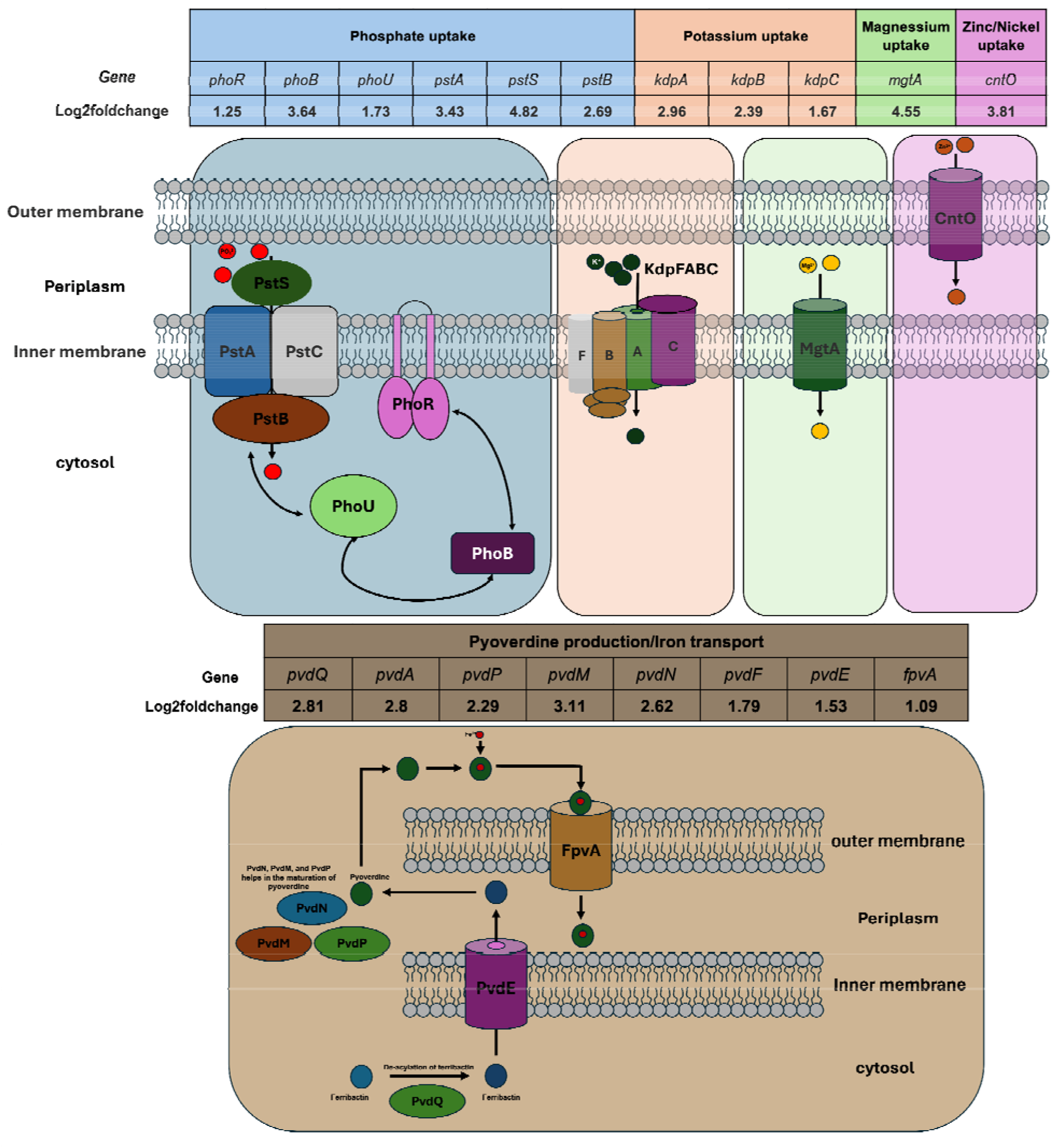
A schematic representation of upregulated genes associated with transmembrane ion transport in *P. aeruginosa* and their roles in ion uptake during ion starvation.

Most of the significantly downregulated genes belong to the arginine and proline metabolism, ribosome, and metabolic pathways KEGG groups (Fig 4). GO group associated with type IV pilus assembly was downregulated (Fig 4; Supplementary File).

### *P. aeruginosa* showed slower evolution of resistance to tEDTA compared to meropenem in a laboratory selection experiment

We assessed how rapidly planktonic cultures of *P. aeruginosa* evolve resistance to tEDTA under strong selection, using an evolutionary ramp approach. We exposed six populations of PA14 and six populations of LESB58 to subinhibitory concentrations of either tEDTA or meropenem and passaged them to fresh caMHB each day, increasing the treatment concentration twofold based on the highest concentration at which visible growth was observed the previous day. At the end of day 7, PA14 grew in 16× MIC (16 µg/ml) of meropenem while only tolerating 1× MIC (0.25% wt/vol) of tEDTA. LESB58 grew in 8× MIC (128 µg/ml) of meropenem at day 7 while only tolerating 0.5× MIC (0.25% wt/vol) of tEDTA. These patterns were identical across the six populations of bacteria used for each strain/treatment combination (**Supplementary File**). These results indicate that it is more difficult for *P. aeruginosa* to evolve resistance to tEDTA than to meropenem under these conditions (Fig 6).

**Fig 6.**
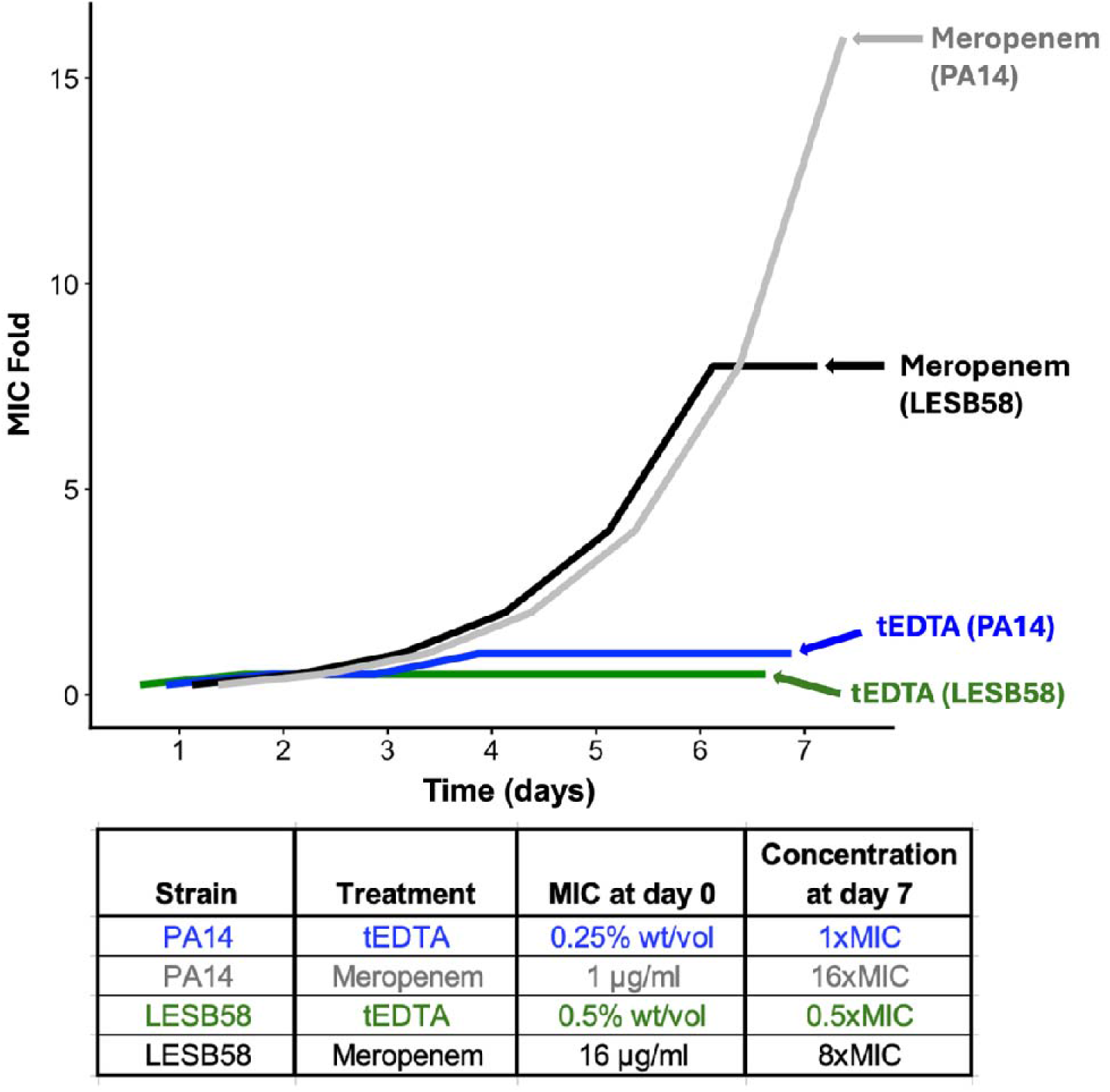
Response of two *P. aeruginosa* strains to selection for tEDTA or meropenem resistance. *P. aeruginosa* PA14 and LESB58 were subjected to subinhibitory concentrations of tEDTA or meropenem and subsequently passaged in different concentrations based on the previous day’s highest concentration. This was done for 7 days. The graph’s axes show the higher of two tested concentrations in which cultures grew at each time point, expressed as a multiple of the ancestral MIC. This experiment was conducted with 6 independent populations for each strain/treatment combination, which all gave the same pattern (**supplementary file**).

### TetrasodiumEDTA showed low cytotoxicity at both inhibitory and biofilm active concentrations

Due to the promising antibacterial and antibiofilm efficacy of tEDTA against different strains of *P. aeruginosa*, we wanted to assess the toxicity of this treatment to human cells. We exposed monolayers of Human Epithelial type 2 (HEp-2) cells to varying concentrations (1%, 0.5%, 0.25%, 0.125%) of tEDTA based on the MIC values in Table 1 and the biofilm active concentrations (concentrations that gave at least 3-log_10_ reduction in PA14 biofilm populations in wound and cystic fibrosis lung models in our previous study (0.5-1%: Orababa et al., 2026)). As a comparator, we have used 16 µg/ml and 1 µg/ml of meropenem as positive controls. These two concentrations represent the meropenem MIC values PA14 and LESB58, respectively. The highest MIC concentration of tEDTA (0.25%) against the *P. aeruginosa* strains showed much lower cytotoxicity effect against HEp-2 cells when compared to the highest MIC of meropenem (16 µg/ml) observed in this study (Fig 7; Supplementary File). Interestingly, biofilm-active concentrations of tEDTA also showed reduced cytotoxic effect compared the concentrations of meropenem tested (Fig 7; Supplementary File). This suggests the potential of tEDTA to be used as a possible treatment for *P. aeruginosa*-associated chronic wound infections.

**Fig 7.**
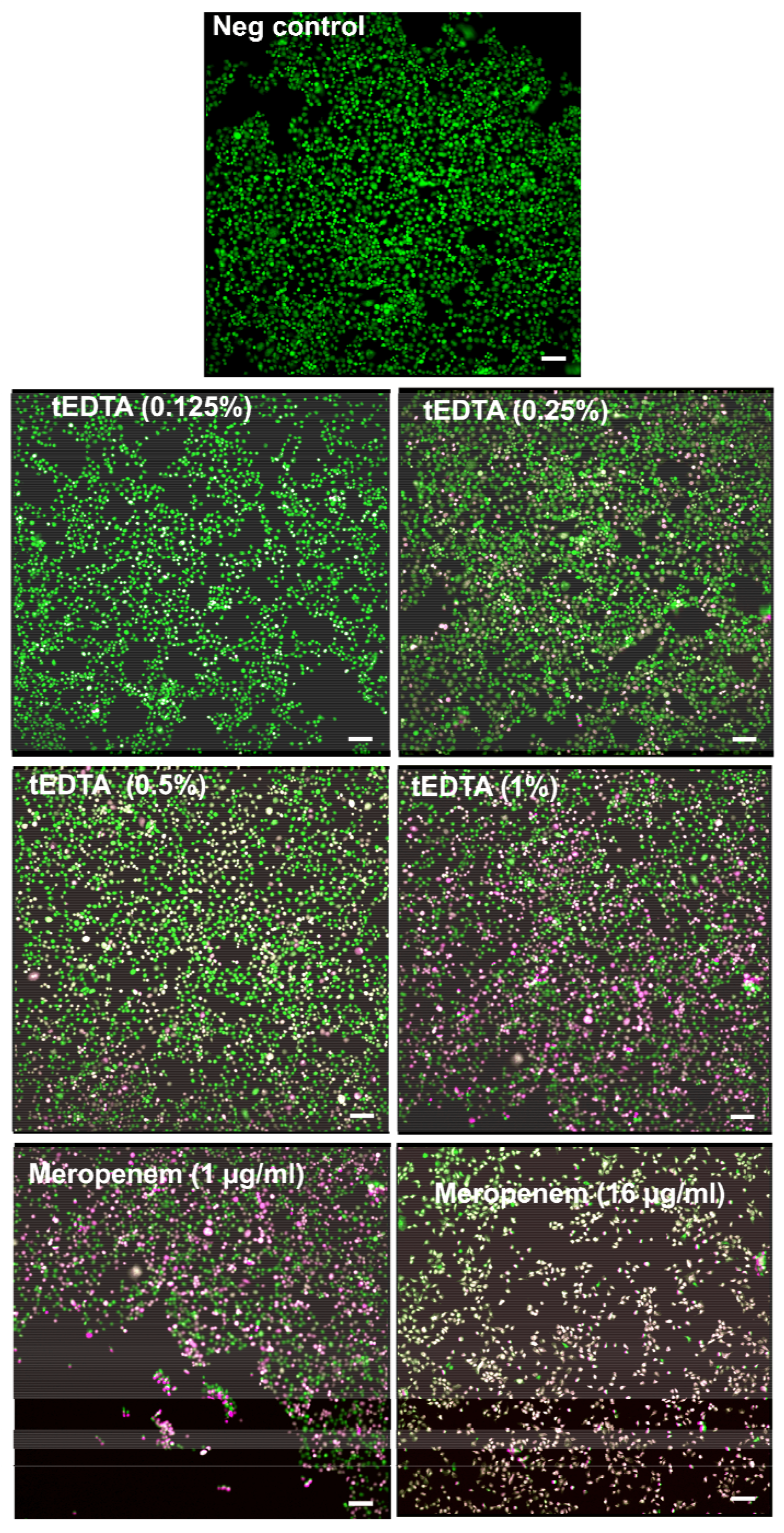
tEDTA has lower cytotoxicity than to meropenem. Human Epithelial type 2 (HEp-2) cells were treated with varying concentrations of tEDTA ranging from 0.125% wt/vol to 1% wt/vol for 2 h and then stained with acridine orange and propidium iodide. Cells were then imaged with Nikon Ti2-E widefield fluorescence microscope (10X objective). Scale bars represent 10 µm.

## DISCUSSION

The growing threat of antimicrobial resistance puts pressure on the need to identify new antimicrobials or repurpose existing compounds to treat drug-resistant infections. Tetrasodium EDTA (4%) is a clinically approved compound in catheter-associated bloodstream infections. We previously identified that tEDTA possesses good antimicrobial efficacy against a lab strain of *P. aeruginosa* (PA14) in both standard and host mimicking media (Orababa et al., 2025). In the present study we reveal that this activity extends to a wider panel of *P. aeruginosa* strains, including clinical strains. We show that regardless of the level of resistance towards meropenem, tEDTA maintains efficacy against both lab and clinical strains of *P. aeruginosa* with MICs of 0.25% wt/vol or less. While tEDTA had the same MIC towards meropenem susceptible (PA14) and resistant (LESB58) strains of *P. aeruginosa*, we identified differences in the growth kinetics of these strains both alone and in the presence of sub-inhibitory concentrations of tEDTA. Our ability to show that the antibacterial efficacy of tEDTA extends beyond laboratory strains and can be replicated carbapenem-resistant strains is one of the strengths of this study.

The current consensus of the mechanism by which EDTA and tEDTA acts against Gram-negatives is by chelating cations in the bacterial membrane resulting in membrane damage (Finegan and Percival, 2015; Moore et al., 2023). However, the mechanism of action of tEDTA is not fully known. For the first time, using membrane disruption assay and transcriptomics, we show that tEDTA disrupts both inner and outer membrane of *P. aeruginosa*, leading to disruption of other cellular processes. Using the membrane sensitive dye, DISC3(5), and membrane impermeable dye, sytox green, we confirmed that tEDTA permeabilises the outer membrane of *P. aeruginosa* PA14, with a rapid quenching of DISC3(5) fluorescence and increased fluorescence of Sytox green. We also showed that there was slow depolarisation of the inner membrane immediately after permeabilization of the outer membrane as evidenced by a slow dequenching of DISC3(5) fluorescence. To the best of our knowledge, this is the first report of the dual membrane activity of tEDTA. This activity is similar to what was previously reported for polymyxin B (Buttress et al., 2022). The membrane permeabilisation activity of tEDTA was further evidenced by the upregulation of genes involved in lipid A and lipopolysaccharide biosynthesis in *P. aeruginosa*.

Our transcriptomics analysis also showed significant enrichment of biological processes associated with phosphate transport, potassium ion transport, inorganic ion transport, and siderophore activity. The upregulation of genes associated with cation transport, including phosphate, potassium and iron is in line with previous report of the ion chelating activity of EDTA (Gray and Wilkinson, 1965; Finegan and Percival, 2015; Moore et al., 2023). Cations are critical for the structural integrity and functional stability of the outer membrane of Gram-negative bacteria: the lipopolysaccharide (LPS) molecules that are present in the outer bacterial membrane are stabilised by ionic bridges formed between adjacent molecules via divalent cations. The chelation of these ions can disrupt these ionic bridges, leading to the release of LPS from the cell surface and a dramatic increase in membrane permeability (Clifton et al,, 2015). This is further evidenced by the upregulation of genes involved in lipid A and lipopolysaccharide biosynthesis. The upregulation of these genes is an indication of the cell attempting to reconstruct the Gram-negative outer membrane in response to the disruption/permeabilization caused by tEDTA. This disruption will facilitate the entry of tEDTA into the cell similar to what was reported for polymyxin B (Savenko et al., 2025).

Cation starvation can also directly inhibit the function of numerous metalloenzymes and cellular systems. For instance, reduced bioavailability of zinc and iron can impede *P. aeruginosa* motility (Li et al., 2026). Interestingly, genes associated with *P. aeruginosa* type IV secretion system (*pilB* and *pilP*) were significantly downregulated. PilB is a cytoplasmic ATPase in *P. aeruginosa* type IV secretion system and has been suggested as an attractive druggable target to treat *P. aeruginosa* infections (Basaran et al., 2025). Future studies should explore the mechanism behind the downregulation of both *pilB* and *pilP* by tEDTA and how this might impact the function of the type IV secretion system in *P. aeruginosa*.

Another interesting *P. aeruginosa* response reported in this study is the upregulation of genes involved in siderophore biosynthesis. Siderophores are secondary metabolites with high affinity for iron (Schalk, 2025). Due to the importance of iron as a co-factor of regulatory proteins and metabolic enzymes, bacterial pathogens devise mechanisms for iron acquisition during infections (Frawler and Fang, 2014). *P. aeruginosa* commonly produces two important siderophores, pyoverdine and pyochelin, that help it scavenge iron for metabolism and colonisation (Jeong et al., 2024. Genes involved in the biosynthesis of both pyoverdine and pyochelin were upregulated in response to tEDTA. This is a possible indication of iron starvation, prompting the cell to produce more siderophores for iron chelation. The iron chelating potential of tEDTA makes it a promising therapeutic option for *P. aeruginosa* infections (Pang et al., 2019).

Carbapenem resistant *P. aeruginosa* is listed as a critical priority pathogen by the WHO. Our MIC results and growth kinetics indicate that tEDTA could be a suitable treatment option towards carbapenem-resistant strains of *P. aeruginosa*. We hypothesised that the impacts on the outer membrane from tEDTA treatment would also result in limited resistance developed towards tEDTA. To the best of our knowledge, experimental evolution has not been previously assessed with tEDTA, especially in carbapenem-resistant *P. aeruginosa*. We performed a ramp style experimental evolution assay with a meropenem susceptible and resistant strain of *P. aeruginosa*. Over a seven-day period, both strains of *P. aeruginosa* (carbapenem-susceptible PA14 and carbapenem-resistant LESB58 strains) rapidly developed resistance towards meropenem. Both PA14 and LESB58 grew in 16x and 8x MIC of meropenem, respectively, at day 7. On the other hand, there was no change in the concentration of tEDTA that supported the growth of both the meropenem-susceptible and - resistant strains of *P. aeruginosa* used in this study. The difficulty of *P. aeruginosa* to evolve resistance to tEDTA is in line with reports indicating that targeting membrane integrity and cellular process display reduced resistance development (Maharramov et al., 2025).

In summary our work shows that tEDTA has good antibacterial efficacy against standard laboratory, clinical, and meropenem-resistant strains of *P. aeruginosa*. We also showed that this antibacterial activity is due to its ability to disrupt bacterial membrane integrity, due to increased expression of genes associated with inorganic ion transport. Importantly our findings reveal that there is limited evolution of resistance towards multiple drug susceptible and resistant strains of *P. aeruginosa*. Together this work highlights the clinical benefits of utilising tEDTA as a therapeutic option to tackle *P. aeruginosa* infections, particularly those exhibiting resistance towards carbapenems such as meropenem. We have also showed that this compound has relatively lower cytotoxicity compared to meropenem and might be a future option in the treatment of drug-resistant infections of *P. aeruginosa* including chronic wound infections.

## METHODS

### Minimum inhibitory concentration assay and growth kinetics

The broth microdilution method was used to determine the minimum inhibitory concentration (MIC) of tEDTA (Sigma-Aldrich, UK) as recommended by the European Committee on Antimicrobial Susceptibility Testing (EUCAST). Briefly, *P. aeruginosa* strains were streaked on Luria-Bertani (LB) agar (Sigma-Aldrich) and incubated at 37°C for 18-24 h to produce distinct colonies. Twice the maximum concentration of tEDTA to be used was prepared in the medium of interest (caMHB (Sigma-Aldrich) and dispensed (100 µl) in the first column of a Corning Costar CLS9018 (Corning Inc., US) 96-well plate. Fifty microlitres of the medium were then dispensed into each of the other wells, after which a two-fold serial dilution was carried out. Bacterial suspensions were prepared by touching 3-4 distinct colonies with a sterile cotton swab and dispersing them in PBS. This was then standardised to a 0.5 MacFarland standard with a spectrophotometer (OD_600_ nm = 0.08-0.10). The standardised bacterial suspension (50 µl) was then inoculated into triplicate wells of each concentration except for the sterile control well (without tEDTA or bacteria). Growth control wells with bacteria (without tEDTA) and with no bacteria were also set up. The 96-well plates (Corning Inc., US) were sealed with Parafilm^TM^ (Bemis) and incubated at 37°C for 18-22 h. The lowest concentration well with no observable growth was taken as the MIC. Experiments were performed with three biological replicates. For growth kinetics, plates were incubated in TECAN Spark at 37°C for 24 h and optical was measured every 15 minutes.

### RNA sequencing and analysis

Overnight cultures of *P. aeruginosa* PA14 were diluted to 0.1 OD_600_ in caMHB, grown to mid-exponential phase, and diluted to 0.8 OD_600_. The *P. aeruginosa* PA14 culture was treated with equal volume (500 µl) of tEDTA or media (control) for 1 h to make a final concentration of 1% wt/vol tEDTA (4×MIC). Treatments were washed off by centrifuging at 10,000 rpm for 10 min. Cell pellets were then resuspended in PBS, centrifuged at 10,000 rpm for 10 min, snap frozen, and sent for RNA extraction and sequencing by Genewiz. Quality of the raw fastq files was checked with fastqc (Leggett *et al.,* 2013), after which trimommatic (Bolger *et al.,* 2014) was used to remove Illumina adapter sequences from the reads. Alignment was done with Bowtie2 (Langmead and Salzberg, 2012). All reads were aligned to *P. aeruginosa* PA14 whole genome sequence (GCF_000404265.1). The aligned sam files were converted to bam format using samtools (Li *et al.,* 2009), after which featureCounts (Liao *et al.,* 2014) was used to obtain read counts. DESeq2 (Love *et al.,* 2014) in R v4.3.3 (R Core Team, 2023) was then used to determine differentially expressed genes, while Gene Ontology (GO) and Kyoto Encyclopaedia of Genes and Genomes (KEGG) analysis were performed with ShinyGO (Ge et al., 2020).

### Membrane disruption (DiSC3) assay

A DiSC3(5) (3,3’-Dipropylthiadicarbocyanine iodide) assay was performed as previously described by Buttress et al. (2022) with a little modification. Briefly, bacterial culture growing at the exponential phase in LB was diluted to 0.5 OD_600_ with PBS containing 0.5 mg/ml bovine serum albumin (BSA) (Sigma-Aldrich, UK) and 2 µM glucose. The addition of BSA was to prevent the binding of DiSC3(5) reagent to the microtitre plate. *P. aeruginosa* cells were washed in PBS containing 0.5 mg/ml bovine serum albumin (BSA) (Sigma-Aldrich, UK) and 2 µM glucose twice and then resuspended. After resuspension, the tube containing washed cells was incubated for 15 min. Bacteria were then transferred into a black polystyrene 96-well plate (Corning Inc., US) and autofluorescence measured (excitation wavelength = 610±10, emission wavelength = 660±10) for 10 min in Tecan SPARK 10 M (Tecan, Switzerland) plate reader, after which DiSC3(5) reagent (Invitrogen) in 1% DMSO was added to give a final DiSC3(5) concentration of 0.5 µM and fluorescence measured for another 10 minutes. Treatments were then added, and the fluorescence was monitored for 1 h. CCCP was used as a positive control.

### Propidium iodide assay

A membrane permeabilisation assay was carried out as previously described by Boix- Lemonche (2020), with modifications. First, bacteria at mid-log phase of growth in LB were diluted to an 0.1 OD_600_ with PBS containing 2 µM glucose, transferred into a black 96-well flat bottom plate CLS3596 (Corning Inc., US) and treated with either tEDTA to final concentrations of 1xMIC and 2xMIC or with water (control). Propidium iodide (Invitrogen) was then added to a final concentration of 5µg/ml and autofluorescence (excitation wavelength = 530±20, emission wavelength = 620±10) was measured for 1 h in Tecan SPARK 10 M (Tecan, Switzerland) plate reader.

### Bacterial cytological staining

Overnight cultures of *E. coli* were grown to mid-exponential phase and diluted to 0.1-0.2 0.1-0.2 OD_600_ in LB. The diluted cultures were then treated with sterile water, 1% wt/vol tEDTA or 2% wt/vol tEDTA for 1 h. Cells were then washed by centrifuging at 10,000 rpm for 5 min and resuspended in PBS. Cells were stained with FM4-64 (red), DAPI (blue) and Sytox Green (green). The stains were then washed, and cells were fixed with formaldehyde. The images were taken with 100x 1 oil immersion objective on a Nikon Widefield Ti2-E (NW1) microscope. Scale bar equivalent to 10 µm.

### Resistance evolution assay

Six independent populations each of *Pseudomonas aeruginosa* PA14 and LESB58 per antimicrobial were started from the ancestor strain. Using an evolutionary ramp approach, populations were serially passaged in caMHB each day for 7 days in the presence of tEDTA or meropenem, starting at a quarter of the MIC and increasing 2-fold each day where visible growth was observed. If no visible growth was observed, the concentration was repeated. This was done for seven consecutive days. Concentration at day 7 was then recorded as a multiple of the MIC of the strain before the start of the evolution experiment.

### Cytotoxicity assay

Human epithelial type 2 (HEp-2) cells (CCL-23 ATCC) were seeded onto 35-mm glass-bottom dishes and cultured in complete Eagle’s Minimum Essential Medium (EMEM) (ATCC) supplemented with 10% (v/v) fetal bovine serum (FBS) (Labtech) at 37°C in a humidified incubator containing 5% CO until approximately 80% confluent. Cells were then treated with tetrasodium ethylenediaminetetraacetic acid (tEDTA) at final concentrations ranging from 0.125% to 1% (wt/vol) for 2 h under standard culture conditions. Untreated cells served as the negative control. Following treatment, cells were incubated with acridine orange (AO; 0.1 µg mL ¹) and propidium iodide (PI; 10 µg mL ¹) prepared in phosphate-buffered saline (PBS) for 30 min at at 37°C, protected from light. Excess stain was gently removed by washing the cells once with PBS before fixation in 4% (wt/vol) formaldehyde for 15 min at room temperature. Following fixation, cells were washed with PBS and imaged. Fluorescence images were acquired using a Nikon Ti2-E widefield fluorescence microscope (10X objective) under identical exposure settings for all treatment groups. Acridine orange-positive cells (green fluorescence) were considered viable, whereas propidium iodide-positive cells (red fluorescence) were considered membrane-compromised or non-viable. Representative images were processed using Omero and ImageJ.

## Acknowledgement

OQO, AK, and CFU were supported by a BBSRC and University of Warwick-funded Midlands Integrative Biosciences Training Partnership (MIBTP) Studentship (BB/T00746X/1). OO was funded by the University of Warwick Institute of Advanced Studies (IAS) and School of Life Sciences, University of Warwick Pump Priming grants. CC and NR are funded by the Medical Research Council (MRC). RG is funded by the Engineering and Physical Sciences Research Council (EPSRC). Human epithelial type 2 (HEp-2) cells (CCL-23 ATCC) were a generous gift from the Nicole C. Robb lab, Warwick Medical School.

## AUTHOR CONTRIBUTIONS

Author contributions following CRedit Taxonomy:

Conceptualisation: OO

Investigation: OO, CC, AK, CFU, SS, LU, BO

Formal analysis: OO, CFU

Methodology: OO, CC, AK, CFU, SS, LU, BO.

Supervision: OO, FH.

Writing – original draft: OO, AK, RG.

Writing – review & editing: All authors.

